# Molecular dynamics simulations of the calmodulin-induced α-helix in the SK2 calcium-gated potassium ion channel

**DOI:** 10.1101/2022.11.02.514819

**Authors:** Rafael Ramis, Óscar R. Ballesteros, Arantza Muguruza-Montero, Sara M-Alicante, Eider Núñez, Álvaro Villarroel, Aritz Leonardo, Aitor Bergara

**Affiliations:** Donostia International Physics Center, 20018 Donostia, Spain; Departamento de Física, Universidad del País Vasco, UPV/EHU, 48940 Leioa, Spain; Centro de Física de Materiales CFM, CSIC-UPV/EHU, 20018 Donostia, Spain; Instituto Biofisika, CSIC-UPV/EHU, 48940 Leioa, Spain

**Keywords:** potassium channel, calmodulin (CaM), secondary structure, protein-protein interaction, molecular dynamics, computational biology

## Abstract

The family of small-conductance (SK) ion channels is composed of four members (SK1, SK2, SK3, and SK4) involved in neuron-firing regulation. The gating of these channels depends on the intracellular Ca^2+^ concentration, and their sensitivity to this ion is provided by calmodulin (CaM). This protein binds to a specific region in SK channels known as the calmodulin-binding domain (CaMBD), an event which is essential for their gating. While CaM-binding domains are typically disordered in the absence of CaM, the SK2 channel subtype displays a small pre-folded α-helical region in its CaMBD even if CaM is not present. This small helix is known to turn into a full α-helix upon CaM binding, although the molecular-level details for this conversion are not fully understood yet. In this work, we offer new insights on this physiologically relevant process by means of enhanced sampling, atomistic Hamiltonian replica exchange molecular dynamics simulations, providing a more detailed understanding of CaM binding to this target. Our results show that CaM is necessary for inducing a full α-helix along the SK2 CaMBD through hydrophobic interactions with V426 and L427. However, it is also necessary that W431 does not compete for these interactions; the role of the small pre-folded α-helix in the SK2 CaMBD would be to stabilize W431 so that this is the case.

## Introduction

Small-conductance potassium (SK) channels are essential for the proper functioning of excitable cells, as they underlie the afterhyperpolarization following the action potential (1–3), and their alterations have been related to several neurological and psychiatric disorders (4). These channels are encoded by four genes known as KCNN1 (SK1), KCNN2 (SK2), KCNN3 (SK3), and KCNN4 (SK4) (5). The SK4 channel is also known as IK (or intermediate-conductance potassium channel) and shows some differences in its sequence, but is structurally analogous to the other three (6). SK channels must be assembled into a tetramer to be functional. Each subunit forming the tetramer is made up of six transmembrane segments (known as S1 to S6) and four helical cytosolic domains (known as hA to hD), as shown in Figure 1A. The transmembrane segments S1 to S4 form the voltage-insensitive domain (ViSD), and S5 and S6 make up the pore domain (PD). The S4-S5 linker is highly conserved among the SK channel family and is relatively long compared to other K^+^ channels. The cytosolic helices hA and hB adopt an antiparallel disposition and run parallel to the plasma membrane, whereas helices hC and hD are important for tetramerization and run perpendicular to the plasma membrane (7).

**Figure 1:**
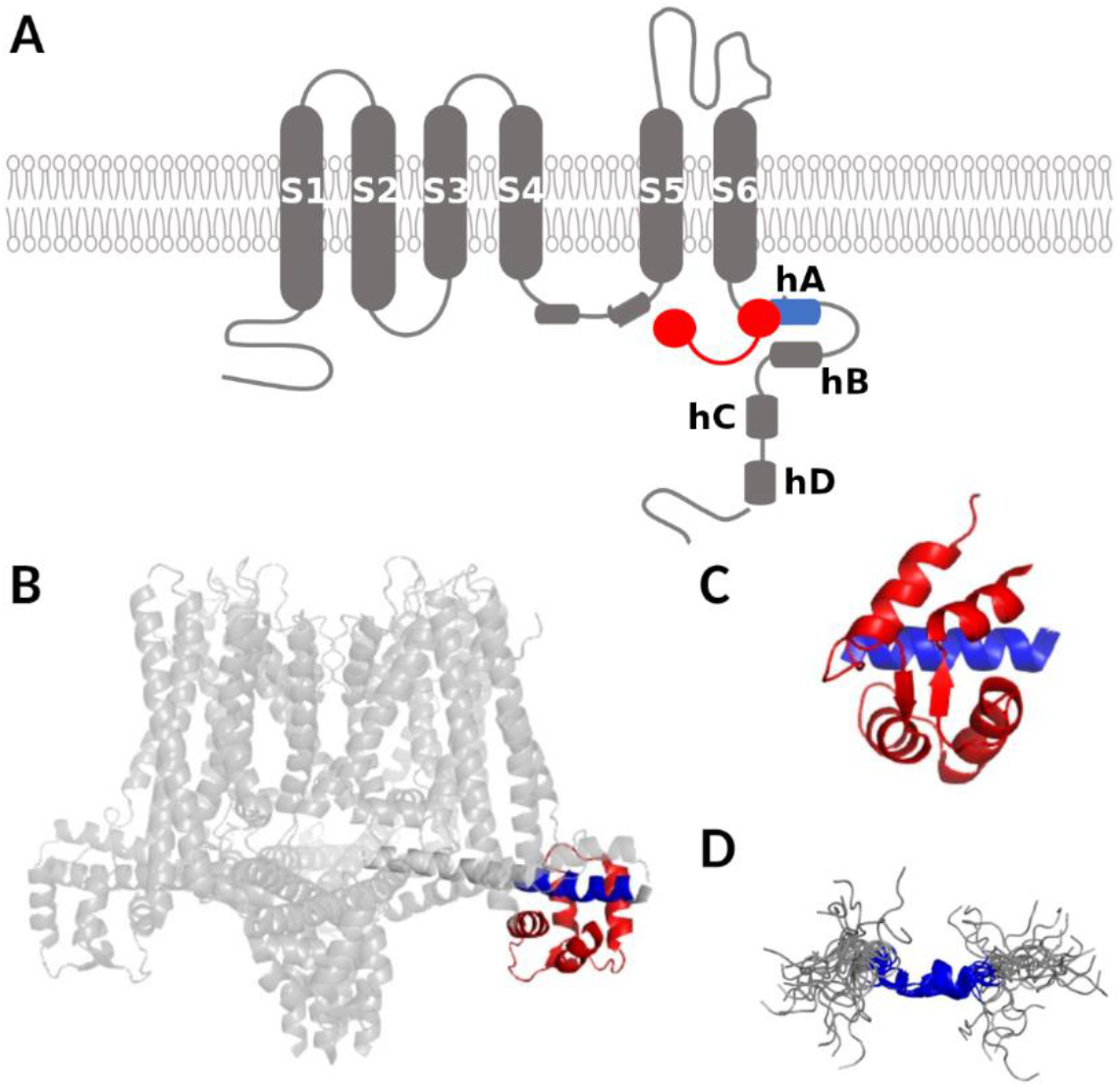
Structural features of SK channels. A: Schematic topological representation of a SK channel subunit (in grey except for helix hA in blue) in complex with CaM (red). B: Cryo-EM structure of the Ca^2+^-free SK4-CaM complex, in which the CaM N-lobe is not visible. The CaM C-lobe is in red, and the helix hA docking region is in blue (PDB: 6CNM (8)). C: Crystallographic structure of the SK2 CaM binding domain (CaMBD) (PDB: 1G4Y (9)) with the C-lobe in red and the helix hA docking region in blue. D: Solution structure of a SK2 peptide with the helix hA sequence, with the binding region colored in blue (PDB: 1KKD (10)).

The gating of SK channels depends on the intracellular concentration of Ca^2+^ ions, which are recognized by the protein calmodulin (CaM). The SK-CaM association is constitutive (i.e. calcium-independent) (11) and the proper binding of CaM to the calmodulin binding domain (CaMBD) of SK channels is crucial for their gating (12). CaM is a 148-residue protein consisting of two globular domains, known as N-lobe and C-lobe, joined by a flexible linker. Each globular domain contains two helix-loop-helix motifs, known as EF hands, each of which can bind one Ca^2+^ ion. In SK channels, CaM binds to hA through its C-lobe, while its N-lobe is not static and is responsible for their pore opening upon Ca^2+^ sensing (13), by engaging to the linker S4a between the S4 and S5 transmembrane domains. This is clearly illustrated in the structure of the full-length CaM-bound SK4 channel, recently resolved by cryo-electron microscopy (PDB ID 6CNM (8)), in which the N-lobe is not visible due to its mobility (Figure 1B). Several other structures of the CaMBD of SK channels bound to CaM (such as that of SK2, Figure 1C) have also been reported (9, 14, 15).

CaM is known for binding a great variety of targets (16) with little in common regarding their amino acid sequences, although most display an α-helical conformation upon binding to CaM through two anchoring hydrophobic residues. However, the details of this recognition process are still uncertain. Both experimental and theoretical approaches have suggested that a mixture of induced fit and conformational selection is needed to explain its promiscuity towards its targets (17, 18). Thus, it is difficult to know *a priori* the mechanistic details of CaM binding to its targets; that is, if it needs a previously folded CaMBD or if it induces the α-helical structure after binding.

The CaMBD of the SK2 channel contains, in aqueous solution and in the absence of CaM, an unusually pre-folded α-helix (10) (Figure 1D) located around a region including a sequence (424-ANVLRETWLIY-434) which resembles the so-called IQ motif, present in multiple CaM targets (19). In a recent work (20), we conducted Hamiltonian replica exchange molecular dynamics simulations which supported the outstanding stability of this pre-folded α-helix. This led us to the hypothesis that pre-helix formation is a requirement for CaM recognition, and that the remaining helix would be induced afterwards.

In this work, we test this hypothesis by conducting Hamiltonian replica exchange molecular dynamics simulations of the SK2 CaMBD bound to the C-lobe of CaM to find out whether or not the pre-helix becomes a full α-helix and, in that case, the role of the different SK2 CaMBD and CaM amino acid residues in this process.

## Results

A short fragment of the CaMBD of the SK2 channel is known to exist in an unusual pre-folded α-helical state in aqueous solution, in the absence of CaM (10). Moreover, in a recent work (20), we could reproduce this pre-folded state through all-atom molecular dynamics simulations on the N421-T437 fragment (the ordered core region of the SK2 CaMBD, shown in Figure 1D). The amino acid sequence of this fragment, together with its NMR-resolved structure (first model in PDB 1KKD (10)), and one of the final snapshots we obtained in our previous simulations, are shown in Figure 2A. In our previous work, we found that the E429-N436 segment was in a stable α-helical conformation while the N421-R428 region remained flexible (Figure 2B). In the presence of CaM, this pre-folded α-helix is known to turn into a fully folded α-helix (14), constituting the hA helix of this ion channel.

**Figure 2:**
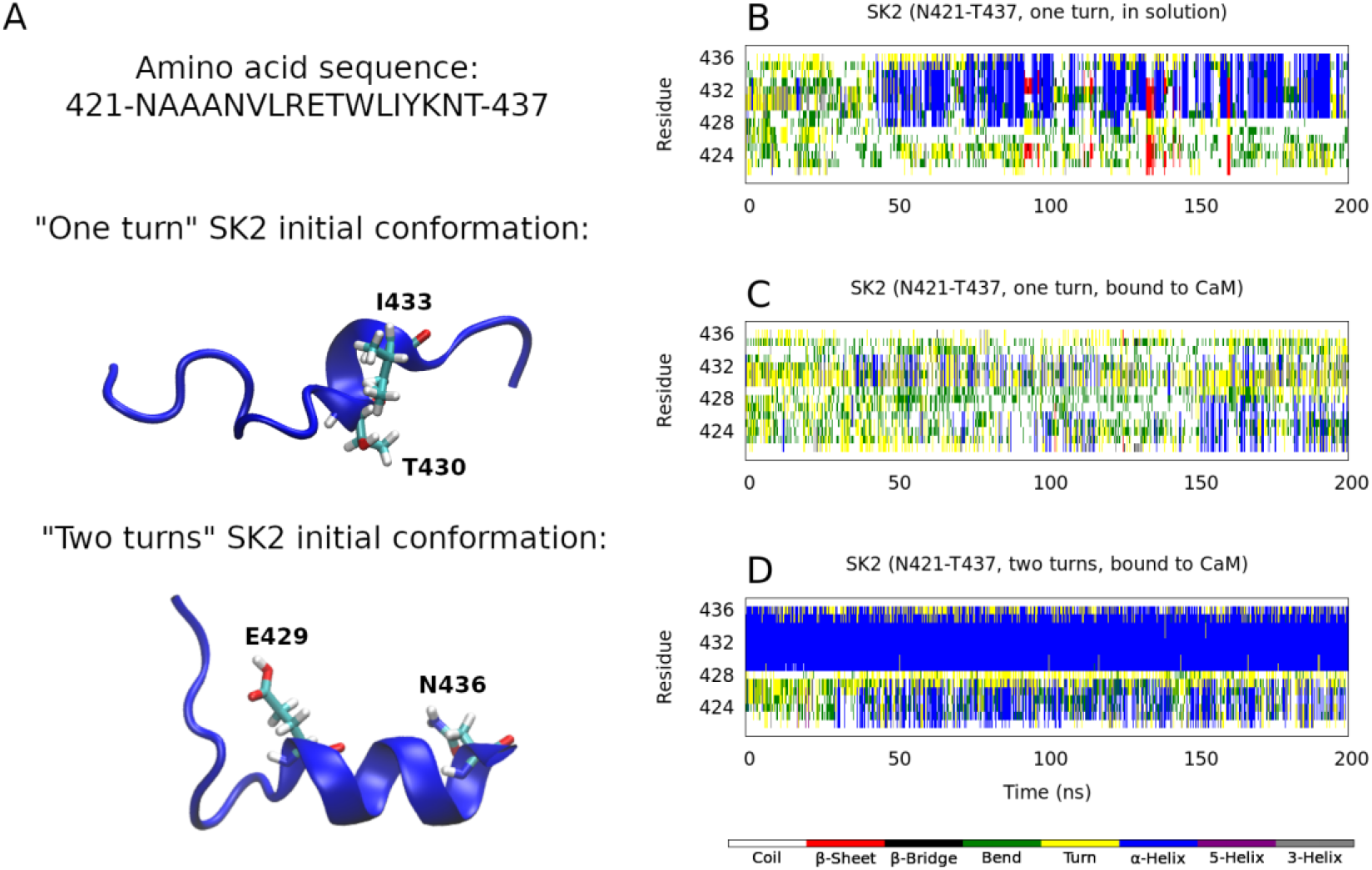
Initial SK2 conformations and secondary structure analysis, according to the DSSP definition (21), calculated along a Hamiltonian replica exchange molecular dynamics trajectory. A: Amino acid sequence of the studied SK2 fragment and cartoon depictions of the 1KKD conformation (10) (with the T430-I433 segment as an α-helix, one turn), and one of the final snapshots in our previous work (20) (with the E429-N436 segment as an α-helix, two turns). B: Secondary structure content in aqueous solution and in the absence of CaM, starting from the 1KKD (“one turn”) conformation. C: Secondary structure content in complex with the CaM C-lobe, starting from the 1KKD (“one turn”) conformation. D: Secondary structure content in complex with the CaM C-lobe, starting from the “two turns” conformation.

In the present work, we conducted all-atom molecular dynamics simulations on the Ca^2+^-bound C-lobe of CaM in complex with the same N421-T437 fragment of the SK2 CaMBD in its pre-folded α-helical conformation. The aim of these simulations was to reproduce the formation of the full α-helix once CaM has recognized and embraced the pre-folded SK2 CaMBD state and analyze the role of CaM in this process.

According to the DSSP (dictionary of secondary structure of proteins) (21), the pre-folded α-helix formed in aqueous solution (N421-T437 sequence of the first NMR model, PDB ID 1KKD (10)) spans just the T430-I433 segment (i.e. it has just one α-helical turn). Our simulations predict that, starting from this conformation, CaM would not induce the formation of a stable full α-helix along the N421-T437 fragment and, moreover, that the pre-folded α-helix would not be maintained (Figure 2C). However, starting from one of the final conformations sampled by our previous simulations in the absence of CaM (20) (with the α-helix spanning the E429-N436 segment, i.e. with two α-helical turns), results not only in the preservation of these two turns, but also in the induction of some α-helical structure on the A422-L427 segment (Figure 2D). This suggests that a certain amount of pre-formed α-helix is needed for CaM to induce a full α-helix. Figure 3 depicts the initial and final snapshots of each simulation.

**Figure 3:**
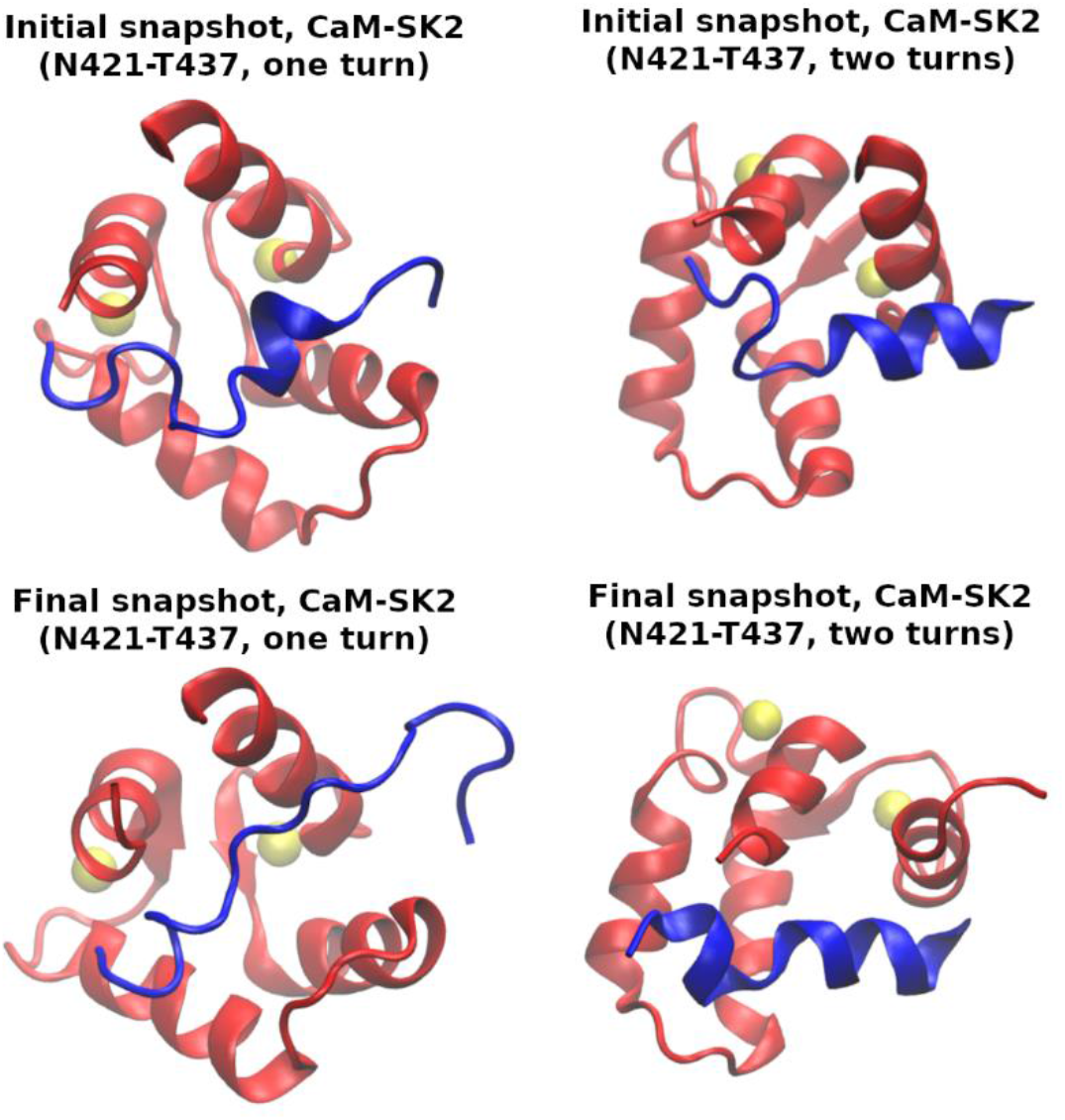
Initial and final simulation snapshots, starting from the 1KKD structure (“one turn”, left) and from one of the final snapshots in our previous work (20) (“two turns”, right). The SK2 calmodulin binding domain rigid fragments (N421-T437) are displayed in blue; the CaM C-lobe, in red; and the Ca^2+^ ions, as yellow van der Waals spheres.

In order to rationalize this result, we analyzed the mobility and solvent exposure of the different SK2 residues in both simulations, expressed as their root-mean square fluctuations (RMSF) and solvent-accessible surface areas (SASA), respectively (Figures 4A and 4B). In the simulation that starts from the “two turns” SK2 conformation (Figure 3, top right), W431 stands out for its much smaller mobility, as its RMSF is almost three times smaller (0.24 nm compared to 0.66 nm). Besides, it shows a notably greater solvent exposure, as its SASA is almost three times greater (1.0 ± 0.3 nm^2^ compared to 0.4 ± 0.3 nm^2^), and it is the only residue for which these (mean ± standard deviation) SASA intervals do not overlap between the “one turn” and the “two turns” systems.

**Figure 4:**
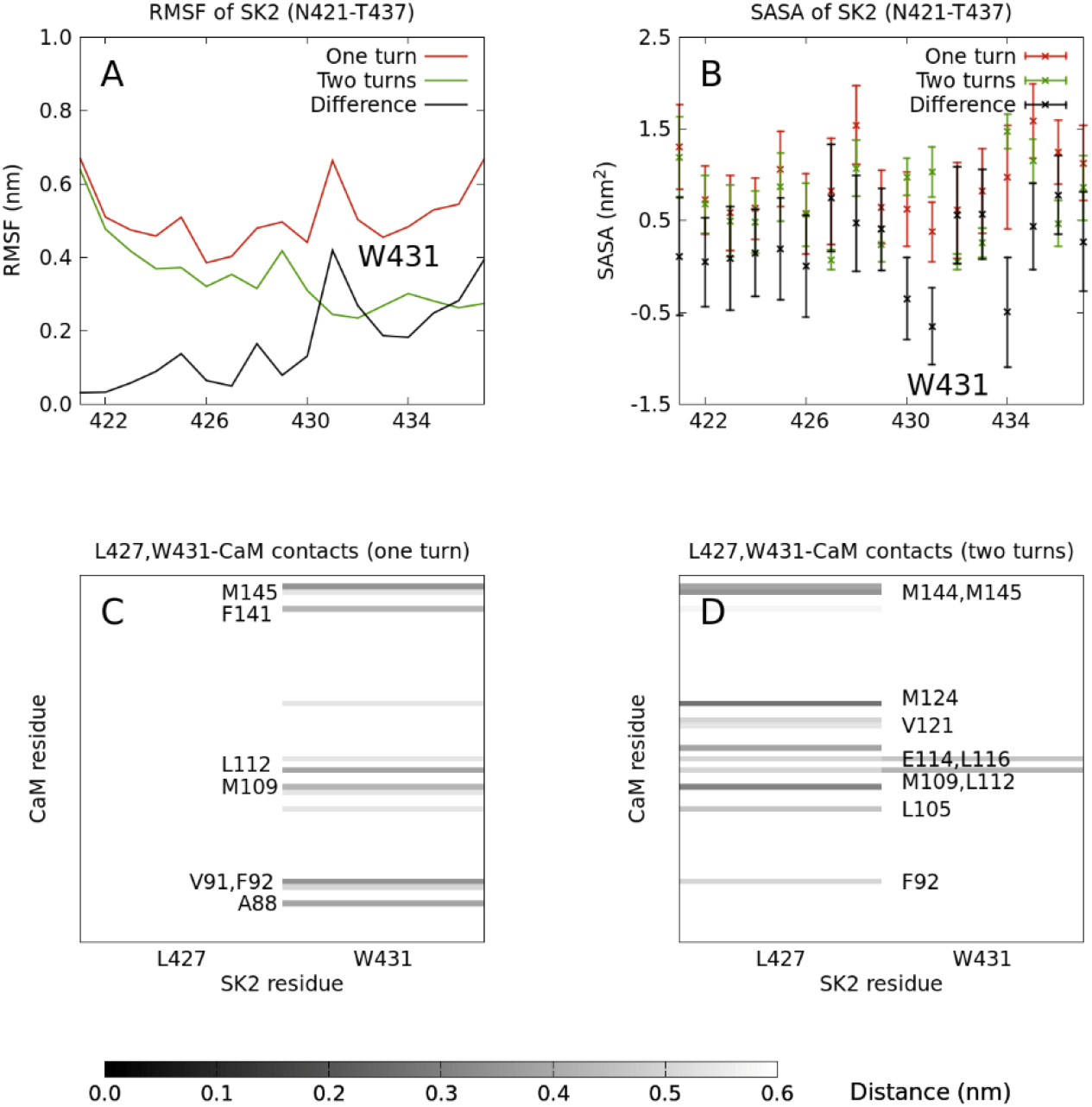
A and B: Root-mean square fluctuation (RMSF), and solvent-accessible surface area (SASA), respectively, of each residue in the rigid fragment of the SK2 calmodulin binding domain (N421-T437), calculated along a Hamiltonian replica exchange molecular dynamics trajectory starting from either the T430-I433 (“one turn”) or the E429-N436 (“two turns”) segment as an α-helix. C and D: Contacts between L427 or W431 and the CaM C-lobe, calculated along these two trajectories. The indicated CaM residues are those whose average distances to either L427 or W431 are at most 0.5 nm.

These results strongly suggest a connection between the mobility and solvent exposure of W431 and the propensity of the system to form a full α-helix, and from now on we will focus on this particular residue. A visual inspection of the “two turns” trajectory shows that W431 is constrained in the E429-N436 α-helix and oriented towards the solvent along the whole run. In the “one turn” simulation, on the other hand, W431 has much more freedom to interact with CaM.

We next analyzed the contacts that W431 establishes with CaM in both simulations. It turns out that several CaM residues with which W431 interacts in the “one turn” simulation coincide with those that interact with L427 in the “two turns” one (Figures 4C and 4D). L427 is the last residue in the CaM-induced α-helical turn, and its stabilization might facilitate the formation of this α-helix.

In the “one turn” trajectory, CaM residues A88, V91, F92, M109, L112, F141, and M145 are, on average, within 0.5 nm of W431. Meanwhile, in the “two turns” trajectory, CaM residues F92, L105, M109, L112, E114, L116, V121, M124, M144, and M145 are, on average, within 0.5 nm of L427. F92, M109 and M145 are the common CaM residues in these two sets of mostly hydrophobic residues (hydrophobic pockets). Therefore, the fact that CaM fails to induce a full α-helix and that the partial helix is not maintained in the “one turn” simulation can be explained by W431 competing with L427 to interact with those CaM hydrophobic residues. On the other hand, in the “two turns” simulation, the stabilization of W431 and its orientation toward the solvent facilitate the interaction between L427 and the CaM hydrophobic pocket, and the eventual induction of the additional α-helical structure.

Given that the orientation of the W431 side chain seems to be determinant for the induction of this α-helix due to this competition for CaM hydrophobic pockets, we ran two additional simulations starting from the same peptide conformations, but rotated by 180 degrees, so that the W431 side chain faces the solvent in the “one turn” trajectory and CaM in the “two turns” one. With this, we intended to avoid this competition between W431 and L427 for CaM hydrophobic residues and see whether this resulted in a greater α-helix formation. These new initial conformations are illustrated in Figure 5A.

**Figure 5:**
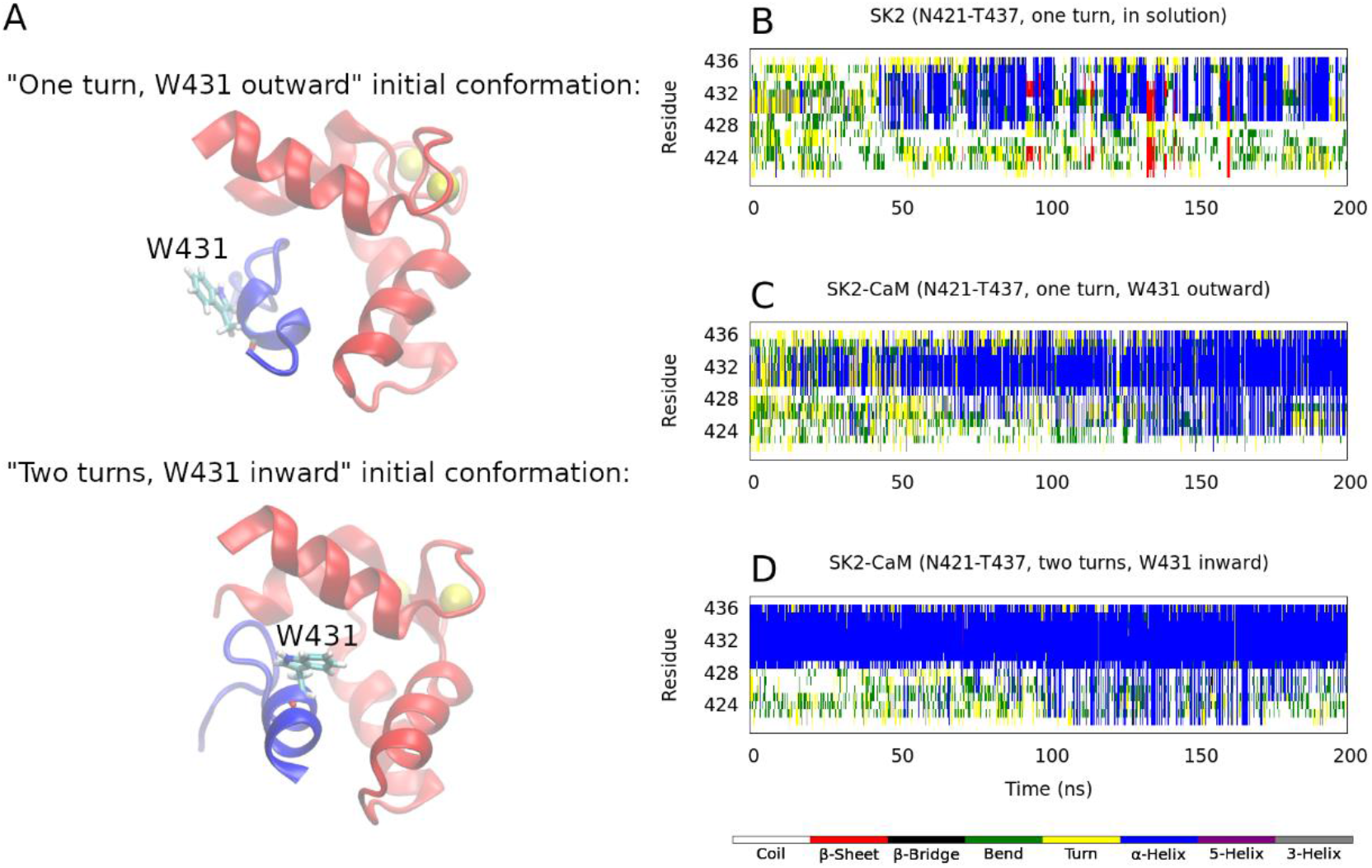
Secondary structure analysis of the simulations starting from two alternative conformations, with SK2 rotated by 180 degrees. A: Starting structure of the “one turn” simulation with W431 pointing outward (i.e. toward the solvent, top) and of the “two turns” simulation with W431 pointing inward (i.e. toward the C-lobe of CaM, bottom). B, C, and D: Same as in Figure 2, but the simulations in the presence of the C-lobe of CaM were started from the conformations shown in A.

As seen in Figure 5C, a much larger degree of α-helical structure can be induced in the “one turn” conformation if W431 is pointing outward (i.e. toward the solvent), in comparison to when it is pointing inward (i.e. toward CaM) (Figure 2C). This α-helix starts forming along the C-terminal end of the peptide (between T430 and L432, i.e. around W431) and then progresses to the rest of it. The “two turns” conformation with W431 pointing toward CaM, however, behaves in a similar way as that with W431 pointing toward the solvent: the E429-N436 α-helix remains stable and some α-helical content is induced along the A422-R428 segment within the second half of the simulation (Figure 5D). Even though W431 is pointing inward, it is stabilized within the pre-helix and not as mobile as in the “one turn” system, facilitating the formation of the full helix. Therefore, the fact that the pre-helix is formed and W431 is stabilized within it is more determinant than its orientation in order for CaM to induce a full α-helix along this SK2 fragment.

The increase in α-helical structure content in the “one turn” system with W431 pointing outward comes with a fair decrease in the RMSF of W431, which drops from 0.66 nm to 0.51 nm (compare Figures 6A and 4A) and a large increase in its solvent exposure, which rises from (0.4 ± 0.3) nm^2^ to (1.3 ± 0.4) nm^2^ (compare Figures 6B and 4B).

**Figure 6:**
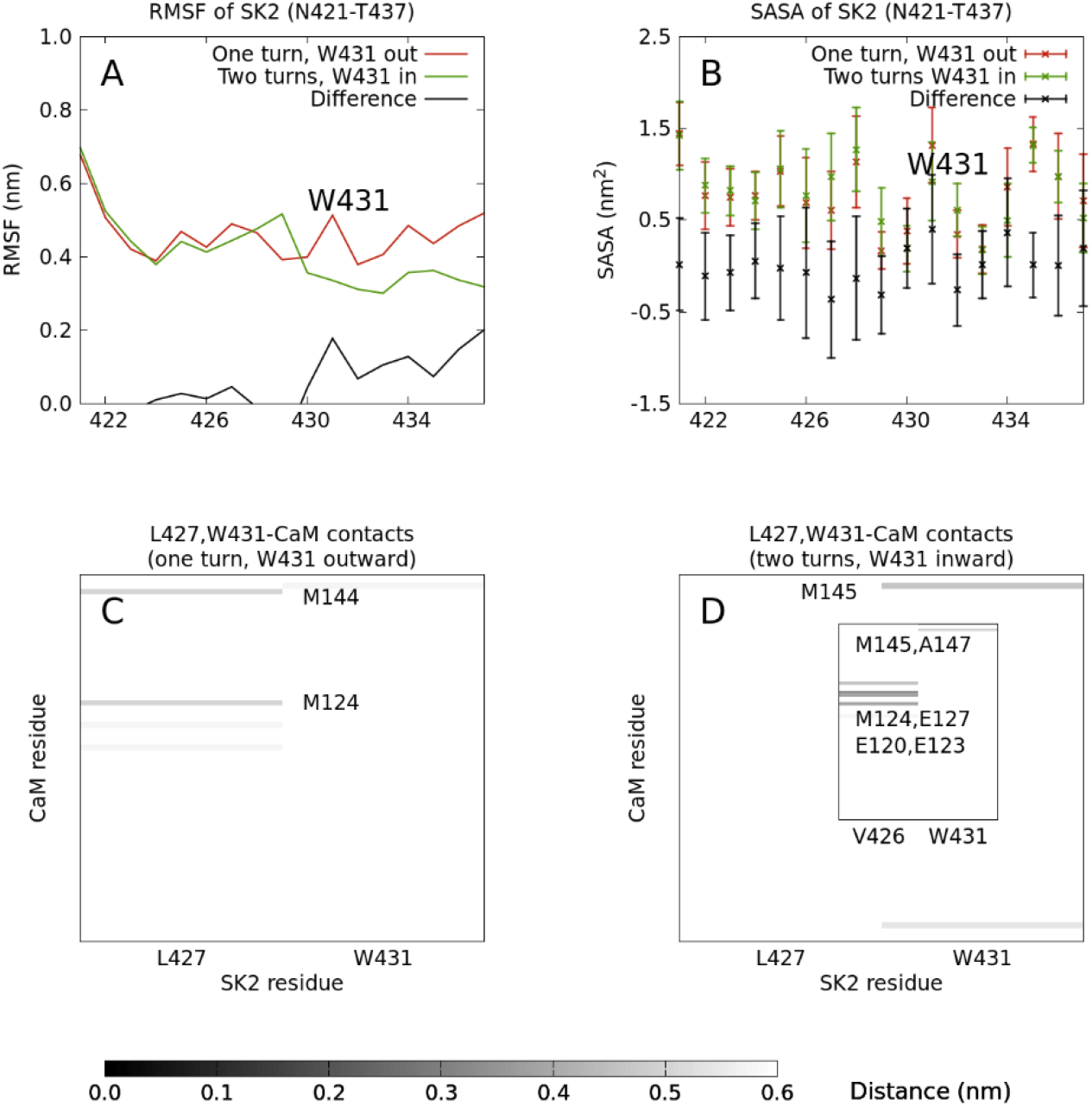
Same as in Figure 4, but the simulations were started with the SK2 fragment rotated by 180 degrees, so that the side chain of W431 was initially pointing toward the opposite direction: outward (i.e. toward the solvent) in the “one turn” system, and inward (i.e. toward the C-lobe of CaM) in the “two turns” system. The inset in D shows the distances between V426 or W431 and CaM C-lobe residues averaged only over the fully helical structures.

An analysis of the distances between either W431 or L427 and CaM (Figures 6C and 6D) reveals a smaller number of contacts in comparison to the other two simulations (i.e., Figures 4C and 4D). In the “one turn” system, W431 does not contact (average distance ≤ 0.5 nm) any CaM residue, which prevents it from competing for the CaM hydrophobic pockets and facilitates the induction of the α-helix. Meanwhile, L427 contacts the hydrophobic CaM residues M124 and M144. In the “two turns” system, on the other hand, W431 just contacts M145 and L427 contacts no CaM residue; an inspection of the trajectory reveals that, in the fully α-helical conformations, it is V426 which is contacting several residues between CaM’s EF3 and EF4, such as E120, E123, M124, and E127 (Figure 6D, inset). Therefore, the interactions of both V426 and L427 would play a role in the induction of α-helical structure along the first half of the studied SK2 fragment.

All in all, in our attempt to describe the formation of a full α-helix along the SK2 CaMBD upon its binding to CaM, we have separately characterized two different states of this system: i) the state where W431 is oriented towards CaM and ii) the state where it is oriented towards the solvent. In the first state, the induction of the full α-helix is less likely than in the second. The maximum effective temperature that we could afford in our Hamiltonian replica exchange simulations (373 K) might not have been enough to observe many transitions between these two states. Therefore, we set up to estimate the energetic barrier separating them.

To do so, we performed metadynamics simulations on a particular angle capturing the relative orientations of W431 and CaM (see the “Experimental procedures” section for details). For both the “one turn” and the “two turns” systems, these simulations yielded two minima separated by a barrier (Figure 7). In the “one turn” case, this barrier is considerably larger than in the “two turns” case (7.6 ± 0.4 kcal/mol compared to 2.4 ± 0.6 kcal/mol). Moreover, while the minimum corresponding to the second state (W431 pointing to the solvent) is close to 2.8 rad in both cases, the minima corresponding to the first state (W431 pointing to CaM) is close to 0.3 rad for the “one turn” system and to 1.2 rad for the “two turns” system. This means that if W431 starts pointing to CaM, it would need to surmount a considerable barrier to point to the solvent, whereas if it starts pointing to the solvent, the barrier would be smaller but it would end up in an intermediate state. The variation of the chosen angle along the trajectories for the “one turn” and the “two turns” systems, and the evolution of the corresponding potential energy surfaces along the last 250 ns of our 2500-ns trajectories, are displayed in Figure S1.

**Figure 7:**
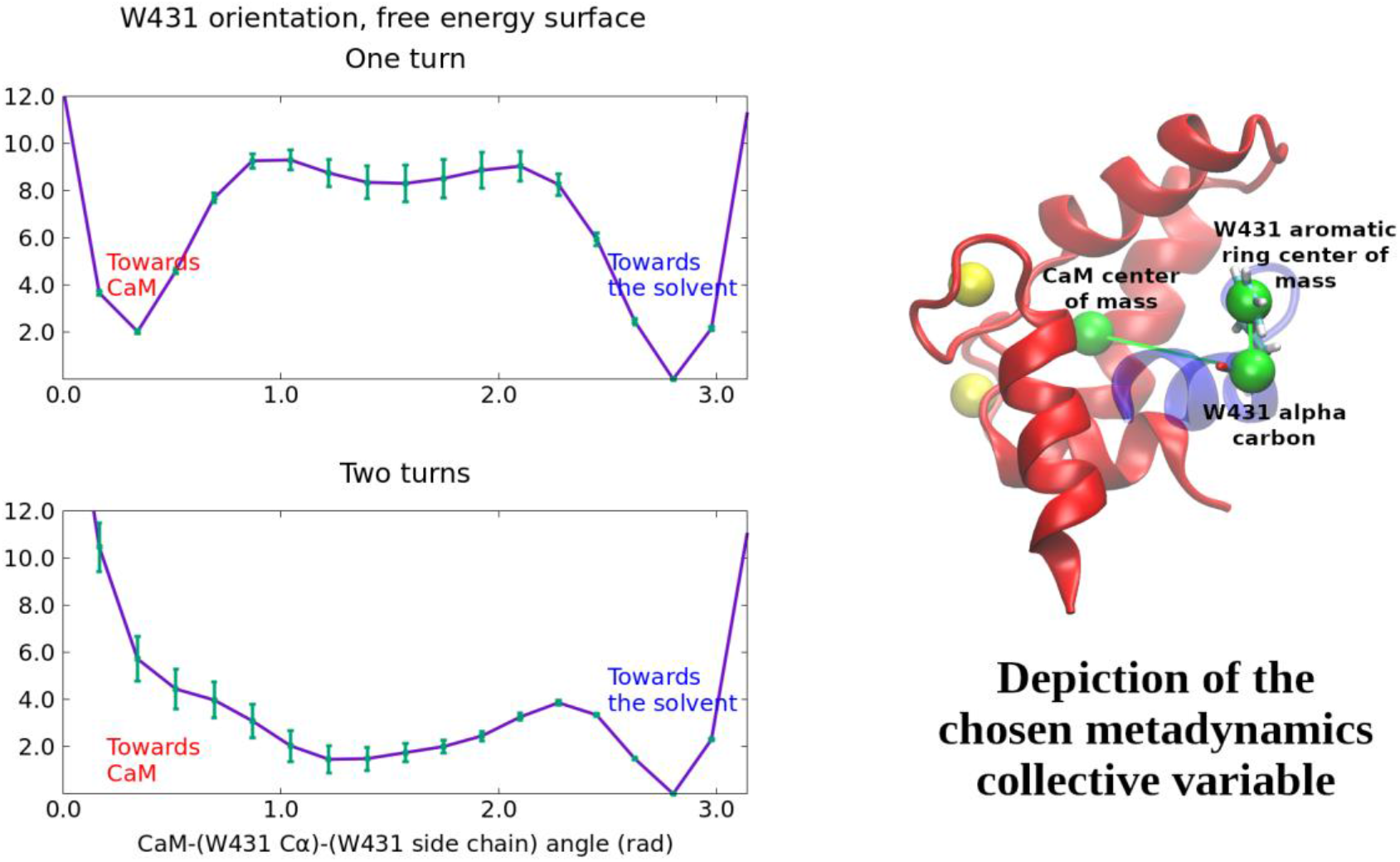
Metadynamics results of the “one turn” (top) and the “two turns” (bottom) systems. Free energy surfaces as a function of the angle between the C-lobe of CaM, the C_α_ of W431, and its aromatic ring (the angle determined by the green spheres on the right panel), along a 2500-ns simulation for each system. The surfaces are the average over the last 250 ns of simulation, and the error bars represent their standard deviation.

## Discussion

The SK2 channel is known to display a peculiar short α-helical region in its CaMBD. This pre-folded region is physiologically relevant, as its disruption precludes CaM binding, yielding non-functional channels (10). In a previous work, we conducted enhanced sampling molecular dynamics simulations to assess the stability of this short α-helix in the absence of CaM (20); the persistence of this helix in time, especially along the E429-N436 segment, is a clear indicator of its stability. In this work, we have applied the same methodology to shed new light into the molecular details of how CaM induces additional α-helical structure along this channel region.

Our simulations have also allowed us to identify the roles played by three particular SK2 residues, namely W431, V426, and L427, and a number of hydrophobic CaM C-lobe residues. V426 and L427 can interact with a hydrophobic pocket including F92, L105, M109, L112, E114, L116, V121, M124, M144, and M145 and favor the formation of additional α-helical structure, whereas W431 may compete for some of these residues (in particular, F92, M109, and M145) and delay the formation of such additional structure. Our results are consistent with X-ray structural studies which identified L427 and W431 as the two residues in the SK2 channel interacting the most with several of the mentioned CaM hydrophobic residues (14), and also with mutagenesis studies which showed that the induction of an α-helix in the SK2 CaMBD was suppressed by the replacement of V426 and L427 with glycine residues (10).

The CaMBD of the SK2 channel is mostly flexible in solution, but transiently samples α-helical structures with up to three turns along the A424-Y434 region (10). The picture emerging from our results is that of CaM *selecting* those conformations where W431 is stabilized within a short α-helix spanning the E429-N436 segment and oriented toward the solvent, which implies that V426 or L427 are pointing toward CaM. Hydrophobic interactions between V426/L427 and CaM would orient the SK2 backbone appropriately for the formation of a full α-helix, whereas those between W431 and CaM would compete with the former and delay the formation of such a helix. Moreover, the fact that W431 is stabilized within the SK2 peptide would compensate for the absence of hydrophobic interactions with CaM.

CaM would then *induce* additional α-helical content along the rest of the N421-T437 SK2 region and, presumably, along the rest of the CaMBD. A schematic depiction of this mechanism is shown in Figure 8. The system would then have to overcome a certain free energy barrier for W431 to point to CaM, since most experimental structures of the CaM-SK2 complex (such as the 3SJQ one (14)) show this orientation. By means of metadynamics simulations, we have estimated a value for this energy barrier lying between 2 and 3 kcal/mol. The rest of the CaMBD or other domains of the channel could help the system to surmount such a barrier. Our metadynamics simulations also suggest that an SK2 conformation without enough pre-formed α-helical content and with W431 pointing to CaM would get stuck in this state (energy barrier between 7 and 8 kcal/mol), thereby delaying the induction of a complete α-helix. The induction of a full α-helix in the SK2 CaMBD from its small pre-folded helix is therefore characterized by a mixture or combination of conformational selection and induced fit mechanisms.

**Figure 8:**
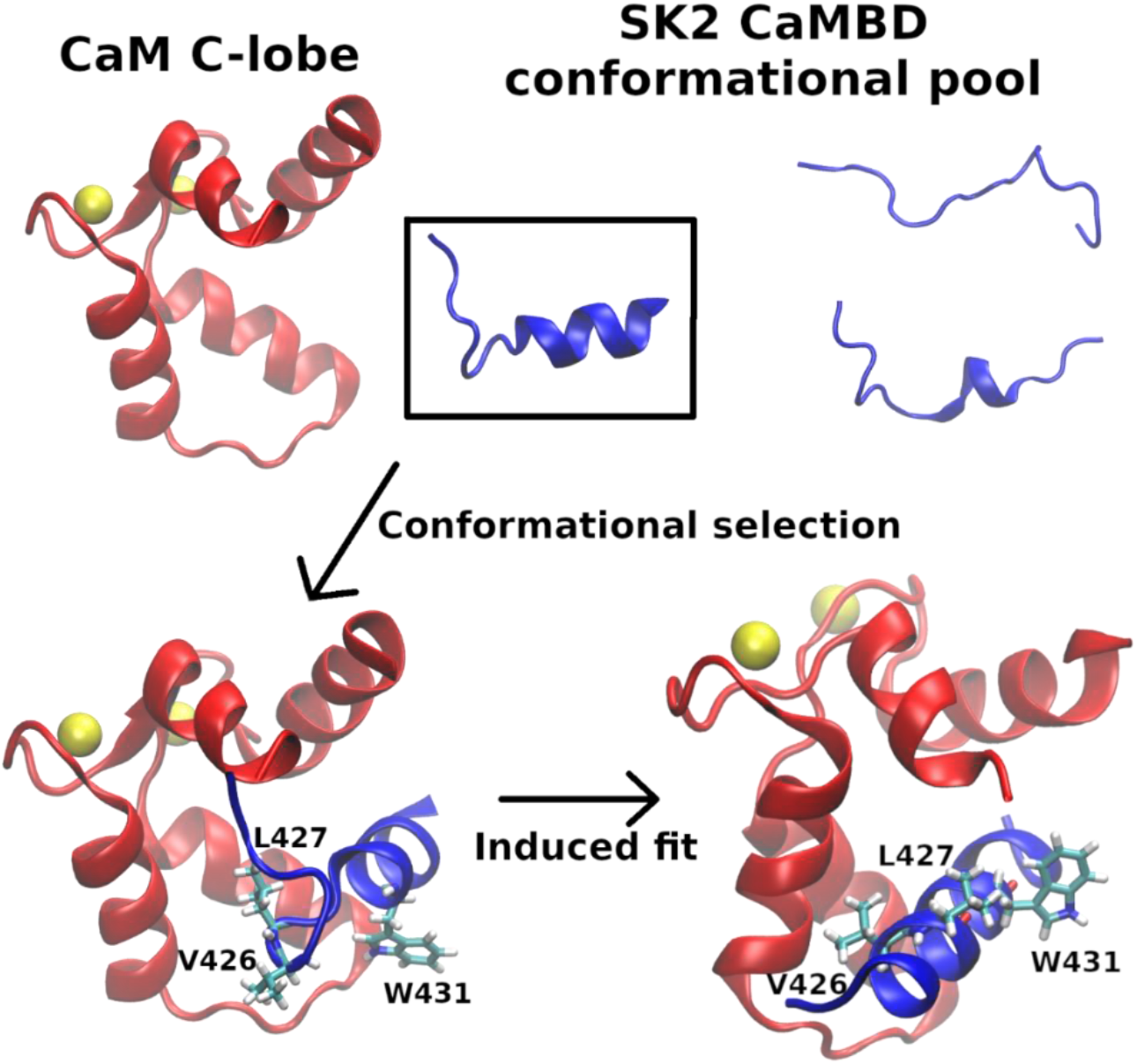
Schematic depiction of the picture emerging from our simulation results. To induce an α-helix along the whole CaMBD of SK2, CaM would *select* those SK2 conformations with at least two α-helical turns, where W431 is less mobile, and then *induce* additional α-helical structure through interactions with V426 and L427.

Residues equivalent to L427 and W431 are present in the CaMBDs of the other SK family members and, more generally, in 24% and 17% of IQ motifs, respectively (20). Figure S2 shows a selection, which includes those of SK1 and SK4 and some related CaM targets. We conducted a secondary structure analysis in the absence of CaM, analogous to that of our previous publication (20), of these CaMBDs (Figures S3 and S4). SK1, SK3 (which are identical in this region), and SK4 contain V, L, and W residues at positions equivalent to those of SK2, and also show a persistent α-helix along this region. On the other hand, IQ motifs such as those of KCNQ1, KCNQ2, MYH7, CAC1C, INVS, or IQGA2, which lack one or more of these three residues, display some transient α-helical structure, although not as persistent as in the case of SK channels. This analysis suggests that a CaM target recognition process similar to that described herein can be generalized to the SK family, and may be present in other IQ targets as well, since they exhibit secondary structure formation when immersed in water.

To complement this analysis, we performed ConSurf (22–26) calculations on the 1KKD sequence to compute the evolutionary rate for each residue in the SK2 CaMBD. The ordered core region simulated in this work (i.e. the N421-T437 fragment) is one of the most conserved regions in the whole CaMBD (together with the A402-M410 fragment). Within the ordered core region, W431 and V426 are two of the best conserved residues (besides A423, A424, and R428). The whole CaMBD sequence colored according to the conservation scores of each residue is shown in Figure S5, and computed evolutionary rates for all residues in the N421-T437 fragment are provided in Table S1. W431 is the only residue occuring at that position in all 300 homologue sequences used for the calculation, while only V, I, or L occur at position 426 and L, V, or M occur at position 427. Again, this analysis provides further evidence of the functional relevance of the three residues identified herein and suggests the relevance of the proposed CaM-assisted helix induction mechanism, even in other similar sequences.

Most CaM binding domains display α-helical structure, but it is not clear whether CaM recognizes them once they are properly folded (conformational selection) or whether it induces their proper folding (induced fit). In this work, we have addressed the case of the pre-folded α-helical core of SK2, which becomes a full α-helix upon binding to CaM. To do so, we have used enhanced sampling Hamiltonian replica exchange molecular dynamics simulations.

Our results, which are consistent with several published experimental and computational studies, suggest that this full α-helix induction takes place by a combination of both mechanisms: CaM is necessary to induce a full α-helix in the N421-T437 SK2 rigid fragment. However, it is not sufficient, since there needs to be some pre-folded α-helical content in aqueous solution (in the absence of CaM). The role of this pre-folded structure would be to reduce the mobility of W431 so that it is less likely to compete for α-helix-stabilizing hydrophobic interactions of V426 or L427 with CaM.

This work represents a step forward in the molecular-level understanding of the physiologically relevant interactions of small-conductance Ca^2+^-activated potassium ion channels with CaM, as it describes a novel role of the latter in the recognition and subsequent folding of SK2 channels, which goes beyond Ca^2+^ signaling. Moreover, given that the residues involved in the recognition are conserved in all members of the SK family, it is reasonable to assume that this mechanism occurs in all of them, and even in other CaM targets with similar sequence.

## Experimental procedures

### Systems preparation

The characteristic pre-folded α-helix of the SK2 CaMBD in the absence of CaM is located along an ordered core region spanning the N421-T437 residues, while the rest of the domain is intrinsically disordered (10). We conducted all-atom Hamiltonian replica exchange molecular dynamics simulations on this ordered region of the channel in complex with the Ca^2+^-loaded C-lobe of CaM. Cartesian coordinates for the channel fragment were taken from two different sources: i) the first model of its NMR-resolved structure (PDB ID 1KKD (10)), which contains a single α-helical turn, and ii) one of the final snapshots of our previous SK2 CaMBD simulation in the absence of CaM (20), which contains two helical turns. Our previous simulation started from the aforementioned first NMR model in 1KKD. Cartesian coordinates for the CaM C-lobe were obtained from the X-ray structure of a variant of SK2 complexed with CaM (PDB ID 3SJQ (14)). The SK2 fragment backbone coordinates were translationally and rotationally fitted to the 3SJQ ones in each case to build the initial structures of the simulated CaM-SK2 complexes. To perform such a fitting, the gmx trjconv tool in GROMACS 2021.3 (27–29) was used, with the -fit rot+trans option. Figure 9 schematically depicts the described workflow.

**Figure 9:**
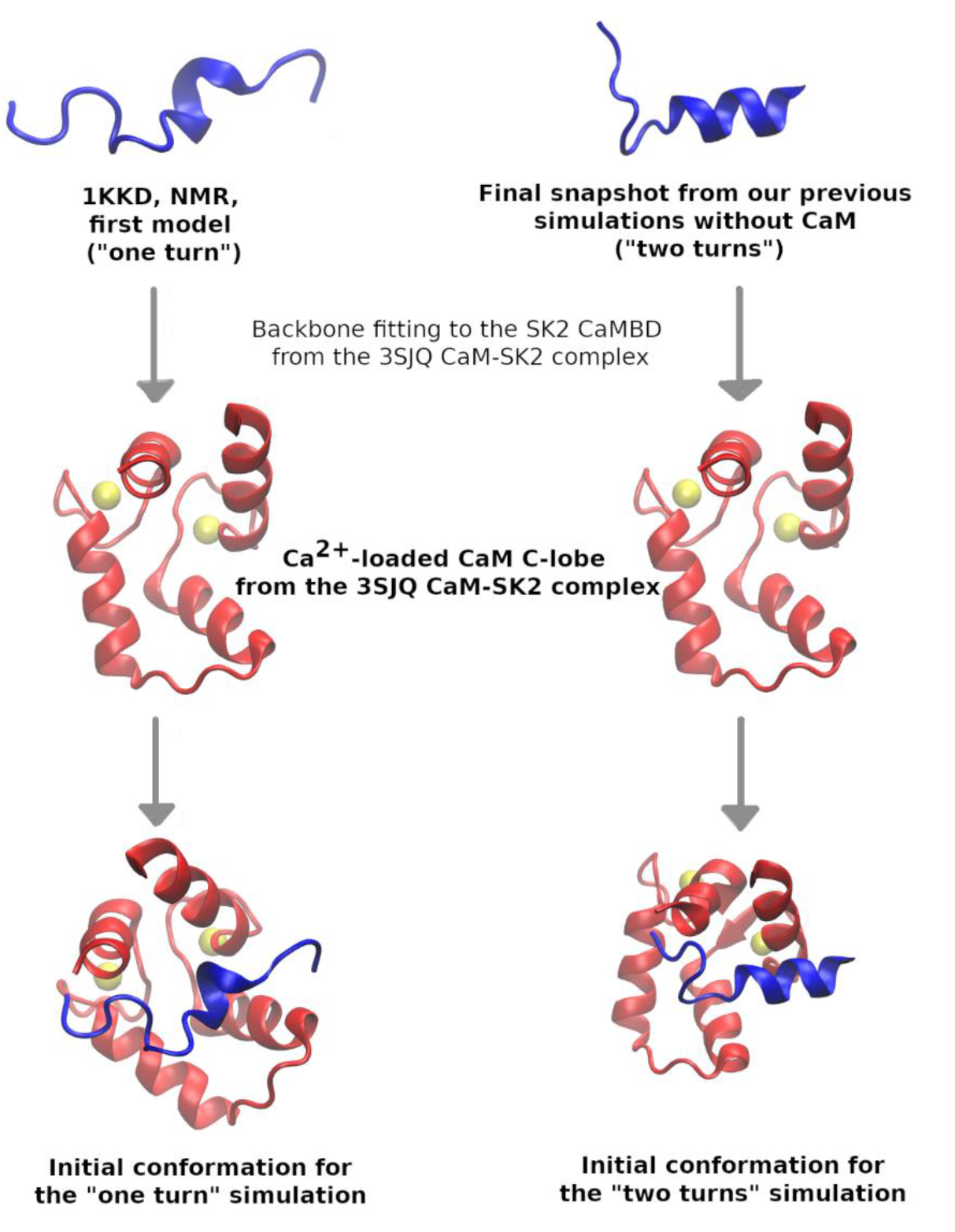
Schematic depiction of the workflow followed to build the initial conformations of the simulated complexes between the SK2 CaMBD and the Ca^2+^-loaded C-lobe of CaM.

### Hamiltonian replica exchange simulations

The CHARMM-36 force field (30) was used for the atoms of the protein and the ions in solution, and the TIP3P force field (31) was used to model the water solvent molecules. In order to enhance the conformational sampling, we employed a variant of Hamiltonian replica exchange molecular dynamics known as replica exchange with solute scaling (REST2) (32). In this variant, all replicas are run at the same physical temperature but on a different Hamiltonian; in particular, a scaled version of the force field used for the first replica. This has the advantage over the standard temperature replica exchange method that we can restrict the scaling to just a part of the system (e.g., the protein atoms, excluding the solvent), so that a much smaller number of replicas is required to achieve a proper sampling enhancement. Information of the simulation will be collected at the first replica, which will follow a trajectory ruled by the usual Hamiltonian but will span high energy conformations, allowing to surmount high energy barriers and enhance conformational sampling.

The part of the system the scaling is restricted to is known as the “hot” region, whereas the rest is known as the “cold” region. The potential energy of the *m*-th replica (*m* ∈ {0,…,*n*-1}), where *n* is the number of replicas) is given by:

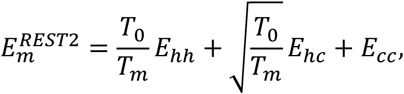

where *E*_*hh*_, *E*_*hc*_, and *E*_*cc*_ are the contributions of the hot-hot, the hot-cold, and the cold-cold interactions, respectively, and *T*_m_ is the “effective temperature” of the *m*-th replica. This last parameter is calculated as:

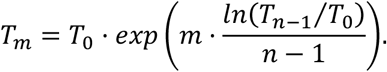

In our simulations, we chose the SK2 fragment as the hot region and all the other atoms as the cold region, as we were mainly interested in the dynamics of SK2. Only the dihedral and the nonbonded terms are scaled, as proposed by Bussi (33), since these are the ones contributing the most to the ruggedness of the potential energy surface. We used a total of *n* = 15 replicas, a reference effective temperature *T*_*0*_ = 298 K and a maximum effective temperature *T*_*14*_ = 373 K. All 15 replicas visited all effective temperatures several times along the 200-ns trajectories, so that there was a proper mixture of replicas, as seen in Figure S6.

### Metadynamics simulations

Our REST2 simulations revealed that the orientation of the W431 side chain in the SK2 CaMBD (toward the C-lobe of CaM or toward the solvent) was relevant for the induction of an α-helix spanning the whole N421-T437 fragment of this domain. For this reason, we decided to perform well-tempered metadynamics simulations on both starting structures to estimate the free energy barrier associated to this orientation change. The chosen collective variable was the angle determined by the center of mass of the C-lobe of CaM, the C_α_ of W431, and the center of mass of the W431 six-member ring. The height of the deposited Gaussians was 2.0 kJ/mol; their width, 0.5 rad; and they were added every ps. The bias factor was equal to 20. These parameters allowed us to cross the relevant barrier in a reasonable simulation time and to explore the two relevant minima, and are similar to those commonly used in biomolecular applications of metadynamics (34–36). A funnel-shaped potential was placed around the SK2 fragment to accelerate the convergence by preventing W431 from exploring conformations too far away from its CaM interaction site.

### Software used

All simulations were conducted with GROMACS 2021.3 (27–29) patched with PLUMED 2.7.2 (37). All systems preparations and analyses were carried out with standard GROMACS and PLUMED tools and in-house bash and awk scripts. All plots were rendered with gnuplot version 5.2 (38) or VMD version 1.9.3 (39).

### Additional details

Once the systems were built, they were placed in a rhombic dodecahedral box with a minimum distance to the walls of 1.5 nm. Water molecules and K^+^ and Cl^-^ ions were subsequently added to achieve electroneutrality and a 0.15 M (physiological) concentration of KCl. Each replica was energy-minimized by means of the steepest descent method until the maximum force was less than 100.0 kJ/(mol·nm). This was followed by a 100-ps NVT equilibration at a physical temperature of 298 K and a 100-ps NPT equilibration at 298 K and a pressure of 1 bar. Both equilibrations were carried out with position restraints on the proteins’ heavy atoms. Production runs were carried out for 200 ns per replica in the NPT ensemble, at the same physical temperature and pressure. Temperature was regulated by means of the velocity-rescaling thermostat (40); and pressure, by the Parrinello-Rahman barostat (41). The leap-frog integrator was used, with a time step of 2 fs. Snapshots were collected every 50 ps, and exchanges between neighboring replicas were attempted every 100 ps. Long-range electrostatics was treated with the PME method (42), and short-range Coulomb and Van der Waals cutoffs were set at 1.0 nm. Periodic boundary conditions and the minimum image convention were employed throughout.

## Data availability

The scripts used to analyze the results, initial structures, and trajectories are available on our group’s GitHub repository (https://github.com/BilbaoComputationalBiophysics/Articles-Supporting_info/tree/CaM_induced_helix_in_SK2).

## Supporting information

This article contains supporting information (20, 22–26, 43).

## Acknowledgments

The authors thank Donostia International Physics Center (DIPC) for providing access to its computational resources.

## Funding and additional information

We acknowledge financial support from the Department of Education, Universities and Research of the Basque Government and the University of the Basque Country (IT1165-19, KK-2020/00110, and IT1707-22), from the Spanish Ministry of Science and Innovation (projects PID2021-128286NB-100, PID2019-105488GB-I00, TED2021-132074B-C32, and RTI2018-097839-B-100) and from FEDER funds.

## Abbreviations and nomenclature

CaM: calmodulin
CaMBD: calmodulin-binding domain
ViSD: voltage-insensitive domain
PD: pore domain
DSSP: dictionary of secondary structure of proteins
RMSF: root mean square fluctuation
SASA: solvent-accessible surface area
REST2: replica exchange with solute scaling
PME: particle mesh Ewald

