## Supplemental Figures and Tables for "Molecular dynamics simulations of the calmodulin-induced α-helix in the SK2 calcium-gated potassium ion channel"

### Supporting Information

### Contents

**Figure S1**: assessment of the convergence of metadynamics.

**Figure S2**: sequence alignment of different CaMBDs to the hA of the SK channel family.

**Figures S3 and S4**: secondary structure analysis of the other SK family members and other CaMBDs showing IQ motifs.

**Figure S5**: Sequence of the SK2 CaMBD, colored according to the conservation score of each residue.

**Figure S6**: Trajectories of the different Hamiltonian replica exchange replicas in the effective temperature space.

**Table S1**: Evolutionary rates for all residues in the SK2 CaMBD ordered core region.

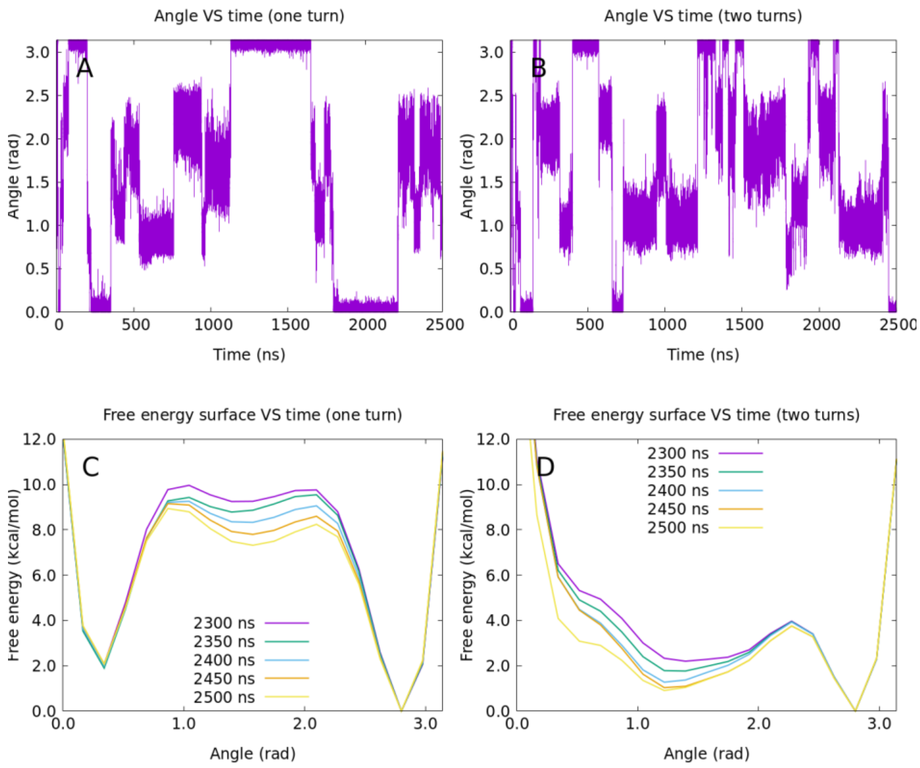


**Figure S1**: Metadynamics convergence. A and B: Time variation of the collective variable (angle between the C-lobe of CaM, the $C_\alpha$ of W431 and its aromatic ring) along a 2500-ns simulation for the "one turn" and the "two turns" systems, respectively. C and D: Potential free energy surfaces projected on this collective variable along the last 250 ns of simulation for the "one turn" and the "two turns" systems, respectively. These surfaces were used to compute the final barrier.

```
sp|KCNN1_HUMAN|Q92952      NAAANVLRETWLIYKHT       17
sp|KCNN2_HUMAN|Q9H2S1      NAAANVLRETWLIYKNT       17
sp|KCNN3_HUMAN|Q9UGI6      NAAANVLRETWLIYKHT       17
sp|KCNN4_HUMAN|O15554      ESAARVLQEAWMFYKHT       17
                           ::**.**:*:*::**:*
sp|KCNQ1_HUMAN|P51787      PAAASLIQTAWRCYAAE       17
sp|KCNQ2_HUMAN|O43526      NPAAGLIQSAWRFYATN       17
sp|MYH7_HUMAN|P12883       SRIITRIQAQSRGVLAR       17
sp|CAC1C_HUMAN|Q13936      FYATFLIQEYFRKFKKR       17
sp|INVS_HUMAN|Q9Y283       DIAAFKIQAVYKGYKVR       17
sp|IQGA2_HUMAN|Q13576      EENVVKIQAFWKGYKQR       17
                                 ::
```

**Figure S2**: Sequence alignment of different CaMBDs to the hA of the SK channel family, carried out with the Clustal Omega tool (1).

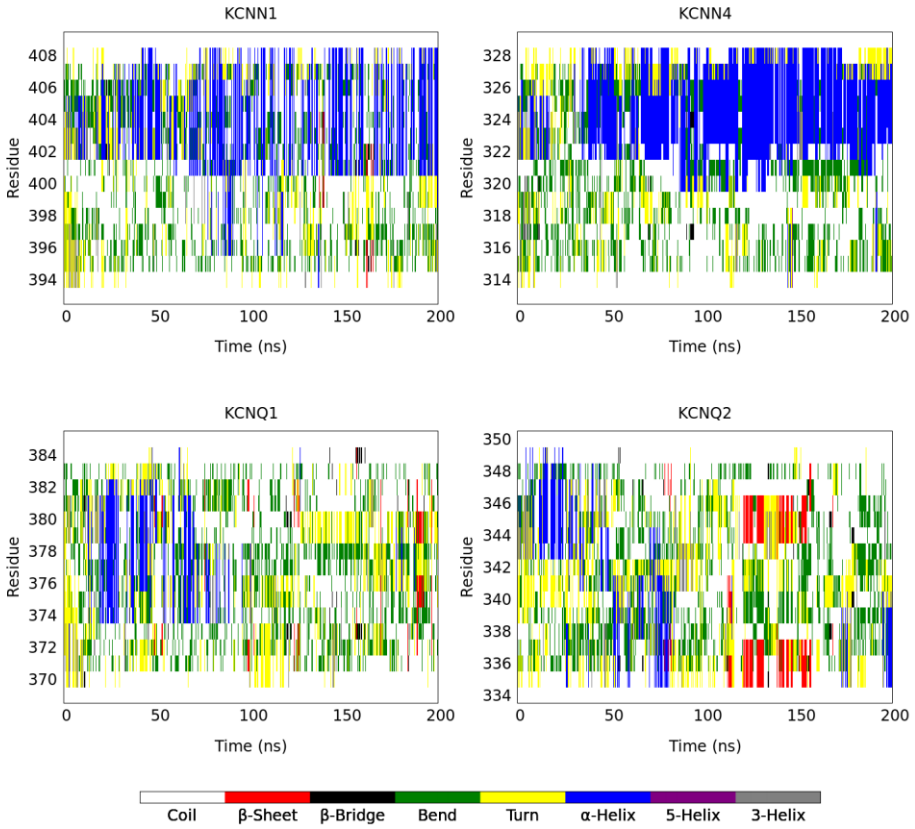


**Figure S3**: Secondary structure content of the other SK channels (KCNN1 (SK1) and KCNN4 (SK4)), as well as the related KCNQ1 and KCNQ2, in aqueous solution in the absence of CaM, calculated as in our previous work (2).

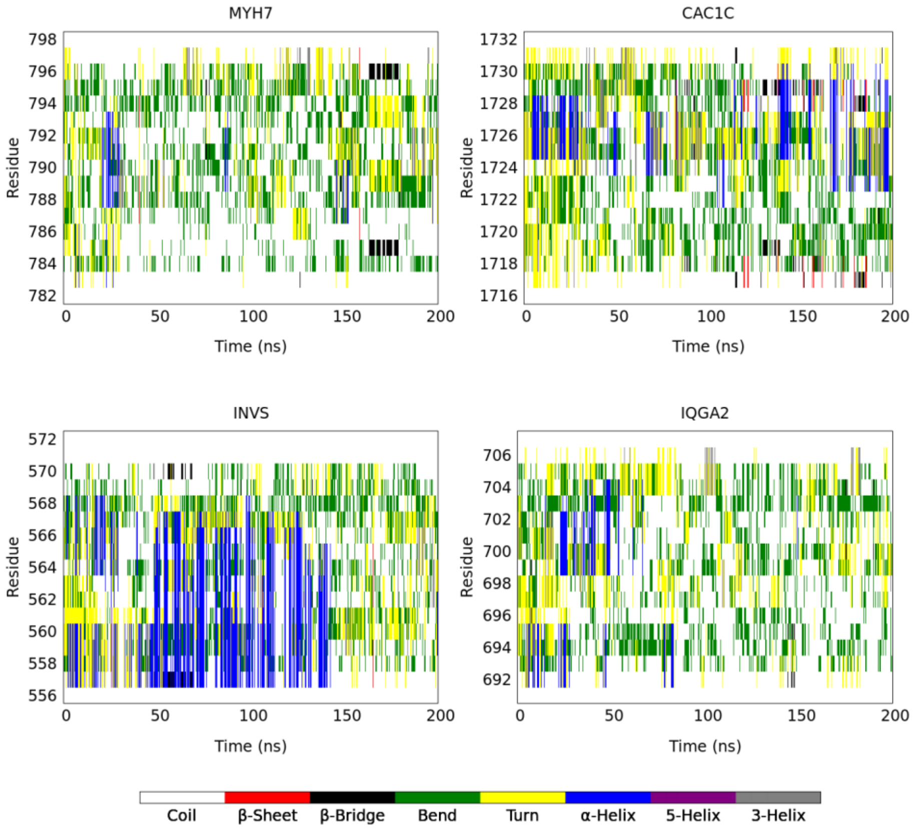


**Figure S4**: Secondary structure content of some additional CaM targets showing IQ motifs: MYH7, CAC1C, INVS, and IQGA2, in aqueous solution in the absence of CaM, calculated as in our previous work (2).

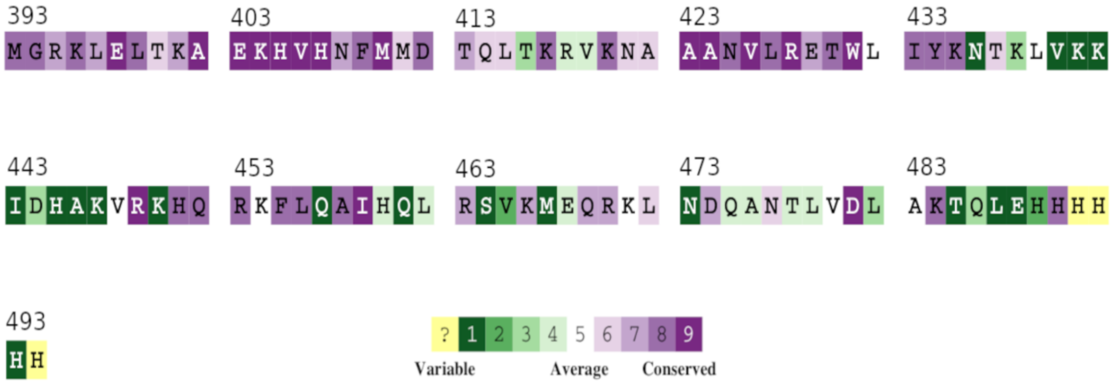


**Figure S5**: Sequence of the SK2 CaMBD, colored according to the conservation score of each residue, calculated using the ConSurf server (https://consurf.tau.ac.il/consurf_index.php) (3–7) with default parameters. The simulated region spans the N421-T437 segment. Residues with the highest conservation scores (in purple) are those with the slowest evolutionary rates (i.e. the most conserved), whereas those with the lowest (in green) are those with the fastest (i.e. the most variable). Residues with unreliable scores due to a high uncertainty are colored in yellow.

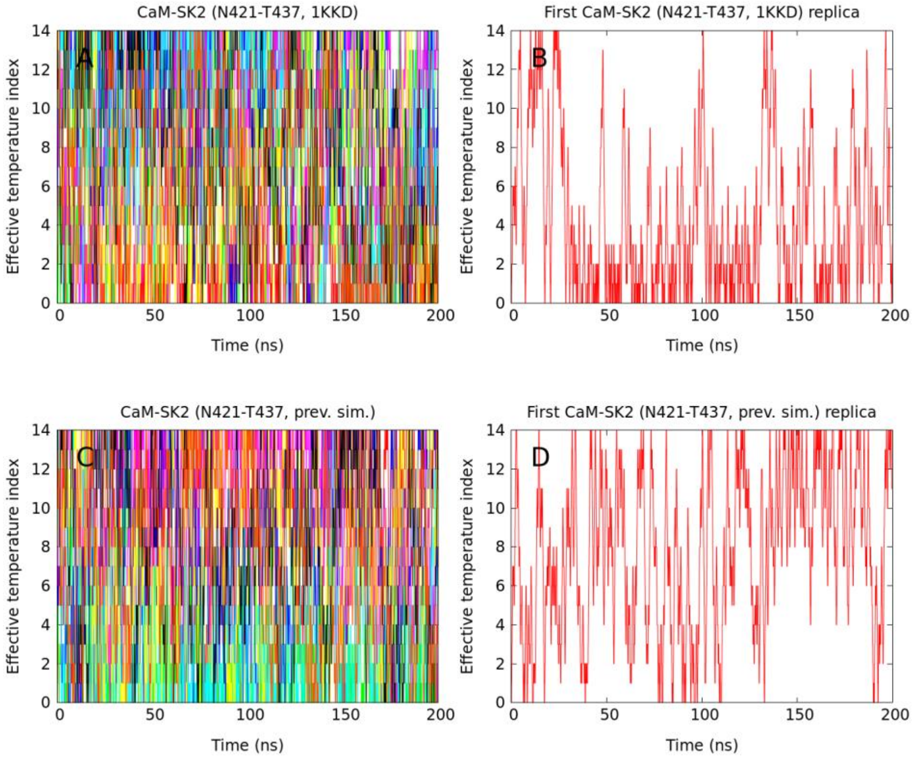


**Figure S6**: Trajectories of all replicas (A and C) and of just the first replica (B and D) in the effective temperature space for the simulations starting from the 1KKD SK2 coordinates (A and B) and from one of the final snapshots in our previous SK2 simulations in the absence of CaM (2) (C and D).

**Table S1**: 50% confidence intervals for the normalized evolutionary rates for all residues in the ordered core region of the SK2 CaMBD (the N421-T437 fragment), relative to the whole CaMBD, as calculated using the ConSurf server (https://consurf.tau.ac.il/consurf_index.php) (3–7) with default parameters.

| Residue | Evolutionary rate 50% confidence interval |
|---|---|
| N421 | (-0.397, -0.043) |
| A422 | (-0.322, -0.043) |
| A423 | (-1.135, -1.078) |
| A424 | (-1.135, -1.078) |
| N425 | (-0.827, -0.643) |
| V426 | (-1.078, -0.969) |
| L427 | (-0.969, -0.786) |
| R428 | (-1.028, -0.903) |
| E429 | (-0.741, -0.530) |
| T430 | (-0.937, -0.786) |
| W431 | (-1.135, -1.028) |
| L432 | (-0.238, -0.208) |
| I433 | (-0.827, -0.643) |
| Y434 | (-0.903, -0.694) |
| K435 | (-0.903, -0.741) |
| N436 | (0.801, 1.643) |
| T437 | (-0.466, -0.147) |
